# Tensor P-Spline Smoothing for Spatial Analysis of Plant Breeding Trials

**DOI:** 10.1101/2021.05.10.443463

**Authors:** Hans-Peter Piepho, Martin P. Boer, Emlyn R. Williams

## Abstract

Large agricultural field trials may display irregular spatial trends that cannot be fully captured by a purely randomization-based analysis. For this reason, paralleling the development of analysis-of-variance procedures for randomized field trials, there is a long history of spatial modelling for field trials, starting with the early work of Papadakis on nearest neighbour analysis, which can be cast in terms of first or second differences among neighbouring plot values. This kind of spatial modelling is amenable to a natural extension using P-splines, as has been demonstrated in recent publications in the field. Here, we consider the P-spline framework, focussing on model options that are easy to implement in linear mixed model packages. Two examples serve to illustrate and evaluate the methods. A key conclusion is that first differences are rather competitive with second differences. A further key observation is that second differences require special attention regarding the representation of the null space of the smooth terms for spatial interaction, and that an unstructured variance-covariance structure is required to ensure invariance to translation and rotation of eigenvectors associated with that null space. We develop a strategy that permits fitting this model with ease, but the approach is more demanding than that needed for fitting models using first differences. Hence, even though in other areas second differences are very commonly used in the application of P-splines, our main conclusion is that with field trials first differences have advantages for routine use.

## 1. Introduction

Designed field trials play a central role in plant breeding and variety testing for most agricultural crops species. The experimental units of such trials (plots) are usually arranged on a rectangular grid and typically show spatial covariance among adjacent plots. There is an ample literature on analysis of field trials using spatial models. One of the earliest papers is Papadakis (1937), who introduced nearest neighbour (NN) methods, an idea taken up later in the seminal papers by Bartlett (1978) and Wilkinson et al. (1983) and subsequently developed further by other authors to cover a wide spectrum of spatial variance-covariance models in common usage today. Spatial models allow making adjustments for irregular spatial heterogeneity and have been shown to improve efficiency in cases where a purely randomization-based mixed model cannot adequately capture the major heterogeneity across the trial area.

One of the earliest papers applying P-splines in field trials, though only in one dimension, is Currie and Durbán (2002). Recently, the use of P-splines (Eilers and Marx, 1996) has been suggested for two-dimensional spatial analysis of field trials (Rodríguez-Álvarez et al., 2018). In this framework, spatial covariance translates into smooth spatial trend. P-splines have a close connection with linear variance and random walk models (Boer et al., 2020). They use B-spline bases for regression on covariates and provide a very flexible framework for smoothing of trends in multiple dimensions, and some of the most prominent applications for this type of modelling are spatial and spatio-temporal smoothing for large environmental datasets (Lee, 2010; Lee and Durbán, 2011; Lee et al., 2013). The simplest approach to spatial smoothing is by geoadditive models (Ruppert et al., 2003, p. 258), where each spatial dimension, i.e. geographical longitude and latitude, is smoothed by separate terms in an additive fashion. Often, however, non-additive extensions are needed to allow for interaction. A key challenge with environmental data is that measurements may be irregularly spaced and hence special care must be taken when modelling the interaction between dimensions. In field trials, matters are considerably simplified by the fact that plots are usually arranged on a regular grid (Appendix A).

Here we will focus on P-spline approaches that allow the partitioning of total variance in an analysis-of-variance (ANOVA) type fashion (Lee and Durbán, 2011; Wood et al., 2013; Wood, 2017), which is also at the heart of the recently proposed P-spline based approaches for field trials (Rodríguez-Álvarez et al., 2018). We seek to use a P-spline framework that is flexible enough for agricultural field trials, yet simple enough for straightforward implementation using mixed model packages. In fact, it is a key feature of P-splines that they have a mixed model representation, meaning that parameters are amenable to estimation by residual maximum likelihood (REML) (Patterson and Thompson, 1971). This makes them particularly appealing because the mixed model framework allows accommodating other features of plant breeding trial data, including genetic correlation among related breeding lines, which can be modelled using genetic marker data (Meuwissen et al., 2001), and random design effects arising from the randomization layout of the trial, as others have pointed out before us (e.g., Verbyla et al., 2018). Also, embedding in a REML framework allows a likelihood-based comparison with other mixed models accounting for spatial correlation.

We consider B-splines with a penalty term of the form *θu^T^D^T^Du*, where *θ* is a penalty parameter, *u* is a vector of regression coefficients with corresponding design matrix *B* and *D* is a differencing matrix (Wood, 2017). In the simplest case, the B-spline basis provides a smooth in just one spatial dimension, but it may also be extended in two dimensions using the Kronecker product of bases for rows and columns (see Appendix A). The penalty determines the smoothness of the coefficients *u* along the spatial dimensions, because it strives for small values of *Du*. The mixed model representation of P-splines rests on the fact that the singular penalty matrix *P* = *D^T^D* is formally equivalent to a precision matrix of a random effect *u*. In order to exploit that equivalence when using a mixed model package for fitting P-splines, *P* needs to be converted to a suitable variance-covariance matrix. In the mixed model representation this leads to a fixed-effects component, representing an unpenalized part, and a random-effects component, representing the penalized part (Currie and Durbán, 2002; Wand and Omerod, 2008; Lee and Durbán, 2011). The resulting models are very similar to those proposed in Verbyla et al. (2018) for smoothing using two covariates based on an integrated squared second derivative penalty used in cubic spline smoothing; but, as will be explained, there are slight differences in the way the unpenalized terms are handled. As our main impetus for this paper is to provide a P-spline framework that is conveniently implemented in a general REML-based mixed model package, we closely follow the philosophy set out in Wood et al. (2013), primarily focussing on such penalties that have just a single parameter and upon conversion give rise to a variance-covariance matrix that is linear in the parameters. It is acknowledged that some penalty matrices that have been proposed for tensor-product P-splines do not have this desirable property and therefore are not considered here in detail. In our discussion, we will briefly review a few important examples of such penalties in order to put our framework into perspective.

A particular focus will be on the choice of differencing matrix *D*. In the context of field trials, the early development of NN methods was based on second differences (Papadakis, 1937; Wilkinson et al., 1983), but later the focus turned to first differences (Besag and Kempton, 1986; Kempton et al., 1994) and the closely related linear variance (LV) model (Williams, 1986), because they are simpler and often provide a good fit. Second differences were rarely used subsequently (but see Green et al., 1985). The recent proposal to use P-splines (Rodríguez-Álvarez et al., 2018) in field trials, however, has led to a revival of this option. In light of the new options that P-splines provide, a revisit of the long-standing question in field trials whether first or second differences are preferable is in order.

This paper has several objectives: (i) To provide a thorough derivation of the two-dimensional P-spline representation as a mixed model, focusing on an ANOVA-type decomposition and paying particular attention to the separation between penalized and unpenalized terms, (ii) to show how such P-splines are intimately related to other commonly used spatial models, such as LV models, (iii) to compare first and second difference penalties in several field trial datasets, and (iv) to compare P-splines empirically with other spatial models such as the first-order autoregressive (AR1) model (Gilmour et al., 1997). The rest of the paper is structured as follows. Section 2 considers a single column of plots to introduce key concepts and notation. This is extended to smooth marginal (main) effects for rows and columns in Section 3. Section 4 shows how interaction between row and column smooths may be added. The framework will be illustrated using examples in Section 5 and compared to other spatial models. The paper ends with a discussion in Section 6.

## 2. Smoothing along a single column of plots

In this section we will set the stage considering a single column of *k* plots. Key elements of the notation that will be introduced step-by-step, also for use throughout subsequent sections, are summarized in Table 1 for ease of reference. Let *B_r_* be a *k* × *m_r_* matrix of *q*-th degree [(*q* +1)-th order] B-spline bases (Eilers and Marx, 1996) for the row-coordinates *h_r_* of the plots (typically the row numbers). Generally, the first interior knot will be placed at the first plot and the last interior knot at the last plot of the column of plots. Often, the remaining interior knots are placed at the other plots. Alternatively, the remaining interior knots can be placed at larger distances between the first and last plot, which reduces the number of random coefficients and hence the dimension of the mixed model equations. The number of B-spline bases, *m_r_*, will generally be given by the number of interior knots (*i_r_*) plus (*q* –1). Thus, if the interior knots are placed at every plot, we will have *m_r_* = *k* + *q* –1.

**Table 1:** Notation for matrices and vectors used for P-splines

| Matrix<sup>§</sup> | Description | Dimensions<sup>§</sup> |
| --- | --- | --- |
| $h_r$ | Row coordinates of plots | $k \times 1$ |
| $B_r$ | $q$-th degree B-spline bases for $h_r$ | $k \times m_r$ |
| $D_r$ | Differencing matrix | $(m_r - p) \times m_r$ |
| $P_r$ | Penalty matrix; $P_r = D_r^T D_r$ | $m_r \times m_r$ |
| $U_r$ | All eigenvectors of $P_r$ | $m_r \times m_r$ |
| $U_{+r}$ | Eigenvectors of $P_r$ with positive eigenvalues | $m_r \times (m_r - p)$ |
| $U_{0r}$ | Eigenvectors of $P_r$ with zero eigenvalues | $m_r \times p$ |
| $d_r$ | Vector of all eigenvalues of $P_r$ | $m_r \times 1$ |
| $d_{+r}$ | Vector of positive eigenvalues of $P_r$ | $(m_r - p) \times 1$ |
| $i_r$ | Number of interior knots | $1 \times 1$ |
| $m_r$ | Number of B-spline bases<sup>%</sup> | $1 \times 1$ |

§ The subscript $r$ refers to rows. For columns, replace subscript $r$ with $c$ (for columns).
§ $k$ = number of rows; $p$ = order of differences. For columns, replace $k$ with $s$ (the number of columns) and $m_r$ with $m_c$ (the number of B-spline bases used for columns).
% $m_r = i_r + q - 1$ , where $q$ is the degree of the B-spline bases; $m_r = k + q - 1$ when the interior knots are placed at the plots. For columns, replace $m_r$ with $m_c$ (the number of B-spline bases used for columns).

The trend down the column may be modelled by the regression term *B_r_u_r_*, where *u_r_* is a vector of *m_r_* regression coefficients. To penalize these, we employ the (*m_r_ – p*)× *m_r_* matrix *D_r_* of *p*-th differences, from which the penalty term is 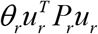, where *θ_r_* is a penalty parameter and 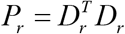 is the penalty matrix. Classical spline smoothing adds this penalty to the residual sum of squares (Green et al., 1985). If a mixed model representation of P-splines is used, the penalty is added to the (residual) log-likelihood (Paul Eilers, in discussion of Verbyla et al., 1999). The penalty is recognized to be equivalent to the term in the loglikelihood for normal random effects, when 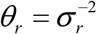, where 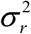 is a variance parameter, and hence the precision matrix is 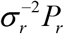. To fit this with a mixed model package, the precision matrix must usually be converted to a variance-covariance matrix. This conversion is not unique because the precision matrix is singular. To resolve this issue, one may use the spectral decomposition 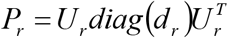, where the columns of *U_r_* are the eigenvectors and *d_r_* is the vector of eigenvalues. Denote the subvector of *d_r_* containing the *m_r_* – *p* positive eigenvalues by *d*_+*r*_ and the corresponding eigenvectors by the columns of *U*_+*r*_. Further, denote the *p* eigenvectors corresponding to the zero eigenvalues in *d_r_* by *U*_0*r*_. Then observing that 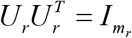, the regression term may be reparameterized by a one-to-one transformation as

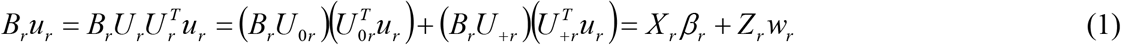

where 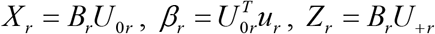, and 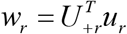. Finally, using the fact that 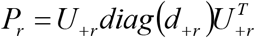 and replacing 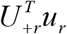 by *w_r_*, the penalty becomes

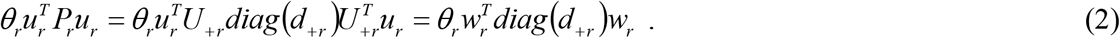

Thus, in the transformed model, only *w_r_* is penalized and is formally equivalent to a random effect, whereas *β_r_*, representing the null space of the penalty, is unpenalized and therefore is formally equivalent to a fixed effect in a mixed model. There are several ways to utilize these key results to fit the P-splines using a mixed model package. In all of these, as a direct consequence of the derivation in (1), it is required to fit the unpenalized fixed-effect term *X_r_β_r_* (Wood et al., 2013; Lee et al., 2021). As regards the random term, we may explicitly fit *Z_r_w_r_* with variance-covariance matrix 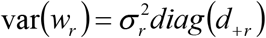. This is equivalent to fitting *B_r_u_r_* as random with singular variance-covariance matrix 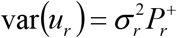, where 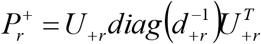 is the Moore-Penrose inverse of *P_r_* (Boer et al., 2020), because 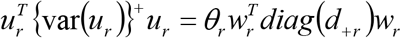 as in (2). In fact, it is proved in Lee et al. (2021) that we may use any generalized inverse of the precision matrix as our variance-covariance matrix and that this provides a unique fit if we simultaneously fit *X_r_β_r_*, representing the null space or unpenalized space, of the penalty matrix. For example, if a positive-definite variance-covariance matrix is required by the mixed model package, one could use the g-inverse 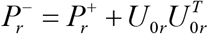. In what follows, we will use the Moore-Penrose inverse 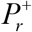, computed from the spectral decomposition, for convenience and refer to a *unique fit* whenever the null space of *P_r_* is fully represented by fixed effects. It is pointed out here that for *p* > 1, *U*_0*r*_, and hence *X_r_*, is determined only up to an orthogonal rotation, i.e. *U*_0*r*_ can be replaced by *U*_0*r*_*R* for any orthogonal rotation matrix *R* (Harville, 1997, p. 538). Statistical packages may differ in the particular form of *U*_0*r*_ obtained in the spectral decomposition of a singular matrix. This indeterminacy is of no consequence, however, because the fit of *X_r_β_r_* by fixed effects, as well as the residual likelihood for the random effects, is invariant to the rotation *R*. In later sections, we will consider two-dimensional extensions of such models that fit specific components of *X_r_β_r_* by random effects for smoothing and in this case the fit is not rotation invariant.

We note in passing that in addition to the smooth plot effect *B_r_u_r_*, we will generally fit an independently distributed residual vector *e* with variance-covariance matrix 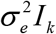, corresponding to a ‘nugget’ variance in geostatistical parlance (Piepho and Williams, 2010). The full mixed model then takes the usual form *y* = *Xβ* + *Zu* + *e*, where the null-space term *X_r_β_r_* is integrated with the fixed effects *Xβ* and the smooth term *B_r_u_r_* is integrated with the random effects *Zu*.

### Special cases

i. When *p* = 1 (first differences), then 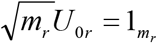 and from the properties of B-spline bases 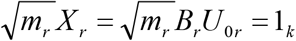, corresponding to the general intercept.
ii. When *p* = 2 (second differences), *X_r_* = *B_r_U*_0*r*_ can be replaced with (1_*k*_⋮*h_k_*), where 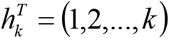, because (1_*k*_⋮*h_k_*) is in the column space of *X_r_* (Lee, 2010; Lee and Durbán, 2011; Lee et al., 2021). Thus, to achieve a unique smooth, we may add a regression on the serial plot number in the fixed part of the model. While the representation of *X_r_* as a regression on plot numbers has some intuitive appeal and is therefore very popular, there is no strict need to use this representation, and one can just use the parameterization in terms of the original eigenvectors, i.e. *X_r_* = *B_r_U*_0*r*_. We think that when it comes to a two-dimensional extension of the smoothing approach, it is in fact preferable to stick with this parameterization (see Section 4.1), because *X_r_* may form part of a term that needs to be smoothed.
iii. If *p* = 1 and *q* = 1, and the interior knots are placed at the plots, we have *m_r_* = *k* + *q* –1 = *k* and *B_r_* = *I_k_*. This model is equivalent to the LV model of Williams (1986) and also to the NN model based on first differences by Besag and Kempton (1986), as well as a first-order random walk model (Boer et al., 2020; Appendix B). Specifically, the spatial variance-covariance matrix of the LV model can be shown to be another generalized inverse of *P_r_* (see Boer et al., 2020).
iv. If *p* = 2 and *q* = 1, and the interior knots are placed at the plots, we have *m_r_* = *k* + *q* –1 = *k* and *B_r_* = *I_k_*. This model is equivalent to the second-differences NN model of Green et al. (1985). For a revealing link of this case with the second-order random walk model (Durbin and Koopman, 2001, p. 39) see Appendix B.

## 3. Smooth marginal effects for rows and columns on a rectangular grid of plots

Now assume that we have plots placed on a rectangular grid of *k* rows and *s* columns (possibly with some missing data). In row-column layouts it is sensible to fit effects for rows and columns. First consider effects for rows. These may be smoothed using random effects. If observations are first ordered by rows and then by columns within rows, this corresponds to fitting random effects *u_r_* with the same specifications for the penalty as given in Section 2 with design matrix *B_r_* ⊗ 1_*s*_. In order to make the smooth unique, we require fitting the corresponding fixed effects *X_r_* ⊗ 1_*s*_. The same kind of effects may be fitted for columns. Thus, we fit random effects *u_c_* with design matrix 1_*k*_ ⊗ *B_c_* and variance-covariance matrix 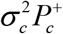. This will be unique if we add fixed effects 1_*k*_ ⊗ *X_c_*, where *X_c_* = *B_c_U*_0*c*_. We may interpret the terms (*B_r_* ⊗ 1_*s*_)*u_r_* and (1_*k*_ ⊗ *B_c_*)*u_c_* as ‘smooth’ marginal effects for rows and columns (Wood 2017, p. 232). When *p* = 1 (first differences), then the two unpenalized terms coincide, i.e. *X_r_* ⊗ 1_*s*_ = 1_*k*_ ⊗ *X_c_* = 1_*k*_ ⊗ 1_*s*_, corresponding to the overall intercept. When *p* = 2 (second differences) the two unpenalized effects have an alternate representation which, apart from the overall intercept, involves a linear regression on row and column numbers with design matrices *h_k_* ⊗ 1_*s*_ and 1_*k*_ ⊗ *h_s_*, respectively. We re-iterate here that while this representation is very popular and is fine for unpenalized terms, we do not advocate this representation for penalized terms as will be needed in the next section.

We note here that when the trial is randomized according to a resolvable row-column design, independent random effects for rows and columns within replicates would be routinely fitted in a randomization-based analysis. Thus, we may regard such a model as a baseline for any spatial extensions, including P-splines. If marginal smooths for rows and columns are added to a randomization-based baseline model, as suggested in this section, they compete with the independent row and column effects. Another way of looking at this is that the independent random effects for rows and columns capture any remaining non-smooth trend not represented by the smooth. Also, the associated variances can be interpreted as nugget for rows and column effects in geostatistical terminology.

## 4. Extending the smooth to cover interaction

Analogous to linear models for two-way analysis of variance (ANOVA), we may augment the smooth marginal effects (*B_r_* ⊗1_*s*_)*u_r_* and (1_*k*_ ⊗ *B_c_*)*u_c_* with a term of the form (*B_r_* ⊗ *B_c_*)*u_rc_*, which will expand the spline space to cover interaction (Wood 2017). There are various choices for the penalty, and special attention needs to be paid to its null space because this may be quite large, e.g., when the penalty is derived from a single Kronecker product of the two marginal penalty matrices *P_r_* and *P_c_*, as considered in Section 4.1. In Section 4.2 we consider a penalty involving a sum of two Kronecker products that is very popular in smoothing (Lee and Durbán, 2011; Wood, 2017; Rodríguez-Álvarez et al., 2018) and is closely related to the intrinsic autoregressive (IAR) model considered in Besag and Higdon (1999).

Before going into details, we would like to stress that our philosophy here is to consider the interaction smooth and its null space as a point of departure. This view is different from other derivations that start with the marginal smooths and then from these derive the interaction smooth terms (e.g., Wood et al., 2013; Wood, 2017; Verbyla et al., 2018). We believe that it is helpful to consider the interaction term in isolation initially because this helps to understand the associated null space and make sure it is fully accounted for in the overall model. As it turns out, in Section 4.1 the null space involves terms that are confounded with the smooth marginal terms introduced in Section 3. This fact makes the derivation a bit involved, but we think it is necessary to take this approach to be clear about the origin and fate of all terms representing the null space of smooth terms for the interaction.

### 4.1. Penalties derived from a single Kronecker product of the two marginal penalties

In this section, we consider penalized differences *D_rc_u_rc_*, where *D_rc_* = *D_r_* ⊗ *D_c_*, and hence the penalty

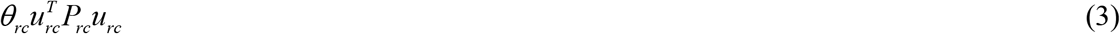

with 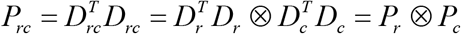, corresponding to random effects *u_rc_* with variance-covariance matrix 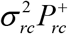, where 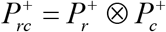. The product-nature of the penalty already hints that its null space is substantial, so substantial in fact, that a smoothing of that null space may be useful, requiring additional penalties, as will be discussed later in this section. To determine the fixed effects needed to represent the null space of *P_rc_*, note that 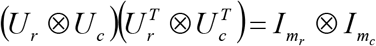 and 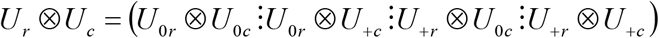, such that using the same approach as in the one-dimensional case [see eq. (1)], the term (*B_r_* ⊗ *B_c_*)*u_rc_* can be transformed one-to-one as

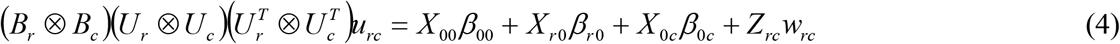

where

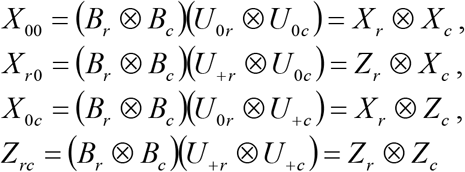

are design matrices, and

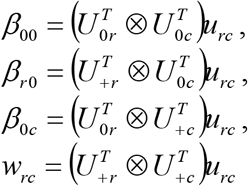

are the corresponding regression coefficients. Analogous to the derivation in Section 2, it emerges that only the term *Z_rc_w_rc_* is penalized because it is associated with the positive eigenvalues *d_+r_* and *d_+c_*, whereas the terms *X*_00_*β*_00_, *X*_*r*0_*β*_*r*0_ and *X*_0*c*_*β*_0*c*_ represent the null space because they are associated with zero eigenvalues and hence need to be fitted as fixed effects to obtain a unique smooth. Note that the dimension of *Z_rc_* is (*m_r_* – *p*)×(*m_c_* – *p*), whereas the null space has dimension *m_r_* ×*m_c_* – (*m_r_* – *p*)× (*m_c_* – *p*), which can be quite large (Wood, 2017, p. 232). This suggests that *p* should be chosen as small as possible.

Importantly, the unpenalized terms *X*_*r*0_*β*_*r*0_ and *X*_0*c*_*β*_0*c*_ will generally show confounding with the marginal smooths introduced in Section 3. It should also be stressed that for *p* > 1 the dimension of the space represented by these terms is larger than that of the marginal terms in Section 3. This is why we think it is crucial in the derivation to initially consider the interaction smooth and its null space in its own right. If we insist that these fixed-effects terms be fitted to ensure uniqueness of the smooth *Z_rc_w_rc_*, then the marginal effect smooths of Section 3 are absorbed into the fixed effects and hence may be dropped. Alternatively, it may be worthwhile to replace the fixed terms *X*_*r*0_*β*_*r*0_ and *X*_0*c*_*β*_0*c*_ by smooth equivalents, which are fitted as random. In this case, however, the smooth *Z_rc_w_rc_* is no longer invariant to the choice of generalized inverse of the penalty matrix *P_rc_* (Lee et al., 2021). Also, because of the confounding, the smooth marginal effects are absorbed by the smooth versions of *X*_*r*0_*β*_*r*0_ and *X*_0*c*_*β*_0*c*_ (and not vice versa!) and hence may also be dropped in this case. It follows that either way, with or without smoothing of *X*_*r* 0_*β*_*r*0_ and *X*_0*c*_*β*_0*c*_, there is no need to explicitly add the marginal-effect smooths described in Section 3; these are accounted for by the terms in the null space of the interaction smooth for (*B_r_* ⊗ *B_c_*)*u_rc_*.

In what follows, we will consider a few important special cases paying particular attention to the null space of (*β_r_* ⊗ *B_c_*)*u_rc_*. and consider how these can be turned into penalized terms. Scrutiny of these special cases then leads to our suggested general approach for tensor-spline smoothing for field trials.

#### 4.1.1 Special cases

i. If *p* = 1, then 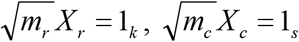 and 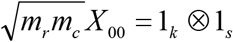, coinciding with the general intercept. The design matrices 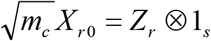 and 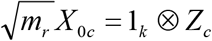 have dimensions (*m_r_* –1) and (*m_c_* –1), respectively, which are usually so substantial that smoothing becomes worthwhile, though this will make the smooth non-unique. For example, for design matrix *X*_*r*0_ we may consider random effects *w*_*r*0_ with variance 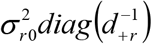, which is equivalent to assuming design matrix *B_r_* ⊗ 1_*s*_ and random effects *u*_*r*0_ with variancecovariance matrix 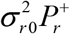. This is recognized as the smooth marginal effect for rows (Section 3). Hence, in this case both *X*_*r*0_ and *X*_0*c*_ may be absorbed into the smooth marginal effects for rows and columns, respectively. Conversely, if we insist that *X*_*r*0_ and *X*_0*c*_ are fitted as fixed, they will absorb the smooth marginal effects for rows and columns, respectively, meaning that the latter can be dropped.
ii. If *p* = 1 and *q* = 1, and interior knots are placed at the plots, we have *m_c_* = *s* + *q* –1 = *s* and *m_r_* = *k* + *q* –1 = *k*. Moreover, *B_c_* = *I_s_* and *B_r_* = *I_k_*. Assuming that the marginal terms *X*_*r*0_*β*_*r*0_ and *X*_0*c*_*β*_0*c*_ are modelled as fixed, this model is equivalent to the LV⊗LV model of Piepho and Williams (2010) and also to the NN model based on first differences along rows and columns by Kempton et al. (1994) (also see Appendix C; Besag and Higdon, 1999; Boer et al., 2020), and all of these models can be viewed as limiting cases of the separable AR1⊗AR1 model (Gilmour et al., 1997; Piepho and Williams, 2010). The terms *X*_00_*β*_00_, *X*_*r*0_*β*_*r*0_ and *X*_0*c*_*β*_0*c*_ correspond to general intercept and row and column effects, respectively. The row and column effects may be many, and it may therefore be worthwhile to fit these as random for recovery of inter-block information (Lee et al., 2021). Again, the corresponding smooth marginal effects in Section 3 may be regarded as representing these random effects already. However, when *X*_*r*0_*β*_*r*0_ and *X*_0*c*_*β*_0*c*_ are replaced by smooths, the smooth *Z_rc_w_rc_* is no longer unique, and also there is no longer equivalence with the LV⊗LV model and twodimensional first differences. Conversely, if we insist on fixed row and column effects to ensure uniqueness, the smooth marginal effects from Section 3 need to be dropped.
iii. If *p* = 2, then *X*_00_ has dimension *p*^2^ = 4 and can be represented by fixed effects for regressions on the row and column numbers and their cross products (Lee and Durbán, 2011). The design matrices *X*_*r*0_ and *X*_0*c*_ have dimensions 2 × (*m_r_* – 2) and (*m_c_* – 2)× 2, and smoothing is worthwhile. For example, *X*_*r*0_ = *Z*, ⊗ *X_c_* could be smoothed separately for the two columns of *X_c_*, which are in the same column space with 1_*s*_. Hence, the smooth would be confounded with the smooth marginal effect for rows. It has random effect vector *w_rc_* with variance-covariance matrix 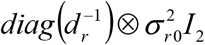, which is equivalent to design matrix *B_r_* ⊗ *X_c_* with random effect *u*_*r*0_ having variance-covariance matrix 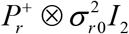. This smooth would represent the smooth marginal effect for rows. It is important to point out here that if *X_c_* was replaced by (1_*s*_⋮*h_s_*), the covariance structure would change, i.e., the model is not invariant to linear transformations with respect to *X_c_*, as is well known for random-coefficient models (Longford, 1993; Wolfinger, 1996). Also note that our suggestion here involves fitting a single penalty for both columns of *X_c_*, rather than two, as is commonly done (Wood et al., 2013; Verbyla et al., 2018). Fitting two separate penalties for *X*_*r*0_ = *Z_r_* ⊗ *X_c_* amounts to fitting the variance-covariance matrix 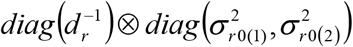 for the random effect *w*_*r*0_, where the variance 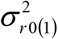 is associated with the first column of *X_c_* and the variance 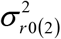 is associated with the second column of *X_c_*. This structure may be compared to a random coefficient model with random intercepts and slopes. In such models, fitting a diagonal variance-covariance for intercepts and slopes means that the model is not invariant to linear transformations of the covariate. This is also the main reason, why we do not recommend replacing *X_c_* by (1_*s*_⋮*h_s_*). Invariance is achieved if we allow a covariance between intercept and slope. Such a covariance will also ensure invariance to a rotation of *U*_0*c*_ and hence of *X_c_*. Thus, we can use the variance-covariance matrix 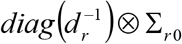, where the two variances on the diagonal of ∑_*r*0_ are 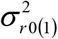 and 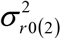 and the covariance is *σ*_*r*0(1,2)_. It is also important to re-iterate that for *p* > 1, the null-space eigenvectors in *U*_*r*0_ and *U*_0*c*_, and hence the matrices *X _r_* and *X_c_*, are only determined up to an arbitrary orthogonal rotation of the eigenvectors (see Appendix D for the special case *p* = 2).
iv. If *p* = 2 and *q* = 1 [a special case of (iii)], the null space can be written in a form involving fixed-effects regressions on row numbers within columns and on column numbers within rows. If these effects are modelled as random for recovery of information, we obviously have a random-coefficient model (Longford, 1993), and this is known to require a covariance among intercept and slope to ensure invariance to linear transformations of the plot coordinates (row and column numbers). We may either model the intercepts and slopes as independent between rows and between columns, or we may use a smooth across the rows and across the columns.

#### 4.1.2 General case

The confounding of the smooth marginal effect introduced in Section 3 and of effects in the null space of *P_rc_* requires adjustment of our notation and actually allows some simplification. The general approach emerging here for ANOVA-type smoothing is to fit fixed effects *β* with design matrix *X_r_* ⊗ *X_c_*, smooth row marginal effects *u_r_* with design matrix *B_r_* ⊗ *X_c_* and variance-covariance matrix 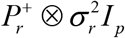, smooth column marginal effects *u_c_* with design matrix *X_r_* ⊗ *B_c_* and variance-covariance matrix 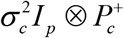, and smooth interaction effects *u_rc_* with design matrix *B_r_* ⊗ *B_c_* and variance-covariance matrix 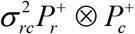. Note that we have re-labelled the effects and variance components here to reflect the absorption of marginal-effect smooths of Section 3 by null-space terms arising from the interaction smooth. We also note that the whole smooth only has three variance components, regardless of the order *p* of differencing, i.e., one for rows, one for columns and one for the interaction. For *p* > 1 there are two further alternatives for smoothing the two marginal effects, which are described here just for the row smooth: In place of 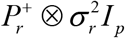 with homogeneous variance for the columns of *X_c_*, we may either use the diagonal structure 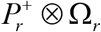 where 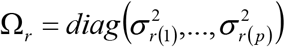 or 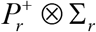, where Σ_*r*_ is an unstructured *p*-dimensional variance-covariance matrix. The former makes the smooth invariant to re-scaling of the spatial coordinates (Wood, 2017, p. 236), but not to translations or rotations (Appendix D). The latter has the advantage of full invariance regarding the representation of *X*_*r*_ and *X_c_*. The lack of invariance to translations or rotations of the diagonal model is particularly relevant, since for *p* > 1 the null-space eigenvectors spanning *X*_*r*_ and *X_c_* are determined only up to an orthogonal rotation. In difference to the smooth terms, the fixed-effects term with design matrix *X_r_* ⊗ *X_c_* is always rotation-invariant for *p* > 1. In summary, for any value of *p*, there are always four terms, i.e., the fixed-effect *X_r_* ⊗ *X_c_*, the two marginal smooths for *B_r_* ⊗ *X_c_* and *X_r_* ⊗ *B_c_*, and the pure interaction smooth for *B_r_* ⊗ *B_c_*.

What was just described as different options for *p* >1 can also be taken to subsume *p* =1 as a special case. For example, for general *p*, the marginal smooth for *B_r_* ⊗ *X_c_* has variancecovariance matrix 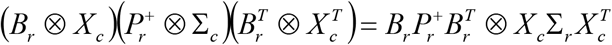. Then for *p* = 1 this reduces to 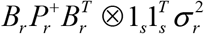. This same reduction also holds for the other two options for *p* > 1, i.e. diagonal and homogeneous. For *q* = 1 this reduces further to 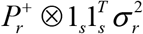, and for knots placed at the plots this corresponds to one of the marginal term for rows of the LV⊗LV model (Piepho and Williams, 2010).

Despite the invariance of the unstructured model to linear transformations of *X*_*r*_ and *X_c_*, the structure is notoriously difficult to fit, as is well known for random-coefficient models in general (Longford, 1993). This is because with poor scaling the variance-covariance matrix ∑*_r_* or ∑*_c_* can have correlations close to the boundary of the parameter space, and it can therefore be difficult to ensure that the matrix remains positive definite. Further problems arise when a variance converges to zero during iterations.

For *p* = 2, we may address these issues by considering an orthogonal rotation of the nullspace eigenvectors. To illustrate this, consider the design matrix *X* = *BU*_0_ (the subscript *r* or *c* is dropped here for simplicity) as needed on the marginal smooth for columns. One possible set of null-space eigenvectors *U*_0_ is (Lee and Durbán, 2011)

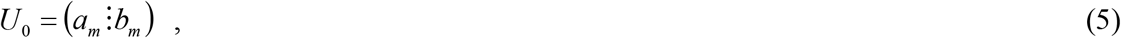

where *a_m_* is an *m*-dimensional vector of 1’s, scaled to have unit length, and *b_m_* is the vector (1,2, …, *m*), centered and also scaled to have unit length. This can be orthogonally rotated to any other permissible set of eigenvectors by *U*_0_*R*(*φ*), where

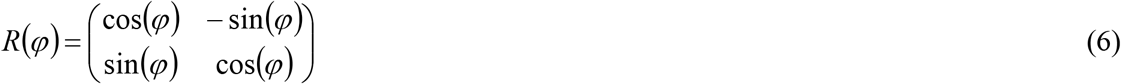

for *φ* ∈ [0,2*π*]. When *U*_0_ is involved in a marginal smooth using the diagonal variance-covariance structure Ω, we may find the optimal rotation by a grid search over *φ*. This is equivalent to fitting a reparameterized version of the unstructured model ∑, with *φ* acting as a third parameter over which the likelihood is profiled. Thus, our proposed procedure for fitting the unstructured model is to perform a grid search identifying a rotation angle that at least nearly maximizes the residual likelihood for the diagonal model, and then use that rotation to fit the unstructured model. The optimization for the diagonal model will ensure that the correlation between the two regression terms associated with the columns of *X_r_* or *X_c_* will be close to zero and safely removed from the boundary of the parameter space. For this purpose, it is not crucial that the correlation is zero exactly, which is why we only need a good guess of the *φ* that would nullify the correlation.

### 4.2. Penalties derived from sum of Kronecker products

Lee and Durbán (2011) consider a penalty of the form

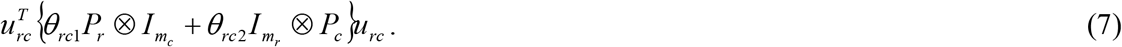

Wood (2017, p. 232) points out that penalties of the form (7) can provide smoother fits than those outlined in Section 4.1. A major advantage of (7) over the interaction smooth in Section 4.1 is that its null space is *X_r_* ⊗ *X_c_* and only has dimension *p*^2^. For *p* = 1, the null space corresponds to just the intercept (Dutta and Mondal, 2015), whereas for *p* = 2 it also comprises the regression on the row and column numbers and their products (Rodríguez-Álvarez et al., 2018). These null spaces can always be accommodated in the fixed-effects part of the mixed model at virtually no cost. A further important consequence of the low dimensionality of the null space is that there is no confounding with the marginal smooths introduced in Section 3, which may be seen as a major advantage of the IAR model. Interestingly, for *p* = 1, *q* = 1, and knots placed at the plots, taking the conditional expectation of *u_i,j_*, the fertility value for the interior plot in the *i*-th row and *j*-th column (i.e., the *i,j*-th element of vector *u_rc_*), given all other *u_i′,j′_*-values, one obtains

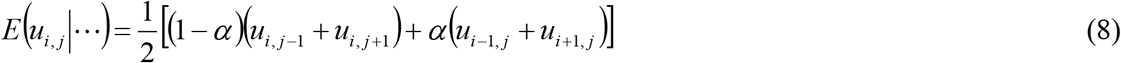

with 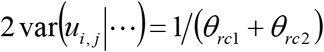, where *α* = *θ*_*rc*2_/(*θ*_*rc*1_ + *θ*_*rc*2_) (modified from Dutta and Mondal, 2015). This is recognized as an NN model, where the central plot is regressed on the nearest row and column neighbours (Julian Besag, in discussion of Bartlett, 1978; Kempton and Howes, 1981). Besag and Higdon (1999) refer to this as the intrinsic autoregressive (IAR) model. Despite this appealing connection with NN models, the penalty (7), as well as variations considered for field trials (McCullagh and Clifford, 2006; Mao et al., 2020), is more difficult to translate to a standard mixed model framework because more than one parameter is involved and hence the inverse of the precision matrix is not linear in the parameters (Wood et al., 2013, p. 345). A convenient option to fit (7) using a mixed model package is by profiling the residual log-likelihood for *α*, i.e., via a grid search over *α* ∈ [0.1] (Besag and Kooperberg, 1995). It must just be kept in mind that when *α* maximizes the residual likelihood at either *α* = 0 or *α* = 1, the penalty in (7) changes to one with a single penalty parameter having a much larger null space, thus sacrificing its desirable properties. This is perhaps the main disadvantage of the IAR model. To obviate the problem at the boundary, a constraint may be imposed such that 0 < *α* < 1, as is done in the Separation of Anisotropic Penalties (SAP) algorithm of Rodríguez-Álvarez et al. (2015). Rodríguez-Álvarez et al. (2018) consider a simplified version of (7),

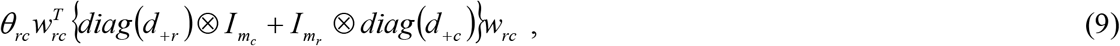

with an associated simplification of (8), which only has a single penalty parameter, allowing this to be fitted easily with a mixed model package. The models implemented in SpATS involve adding marginal smooths for rows and columns with diagonal variance-covariance structures as described in 4.1.

## 5. Examples

### 5.1. A barley trial

Durbán et al. (2003) consider a trial with 252 barley lines laid out as a resolvable row-column design with 2 replicates, 8 rows and 34 columns per replicate [Figure 1(a)]. The replicates were adjacent so that the trial had *k* = 16 rows and *s* = 34 columns in total. We reanalyse this data using several special cases of our P-spline approach. In all models with spatial covariance, we allow the spatial covariance to extend across replicates. For P-splines with both *p* = 1 and *p* = 2, and also for other models, we fit row and column coordinates and their product as fixed regression terms so that the deviance and Akaike information criterion based on the residual likelihood, using twice the number of variance parameters as penalty term, can be used to compare all models (Wolfinger, 1996).

**Figure 1:**
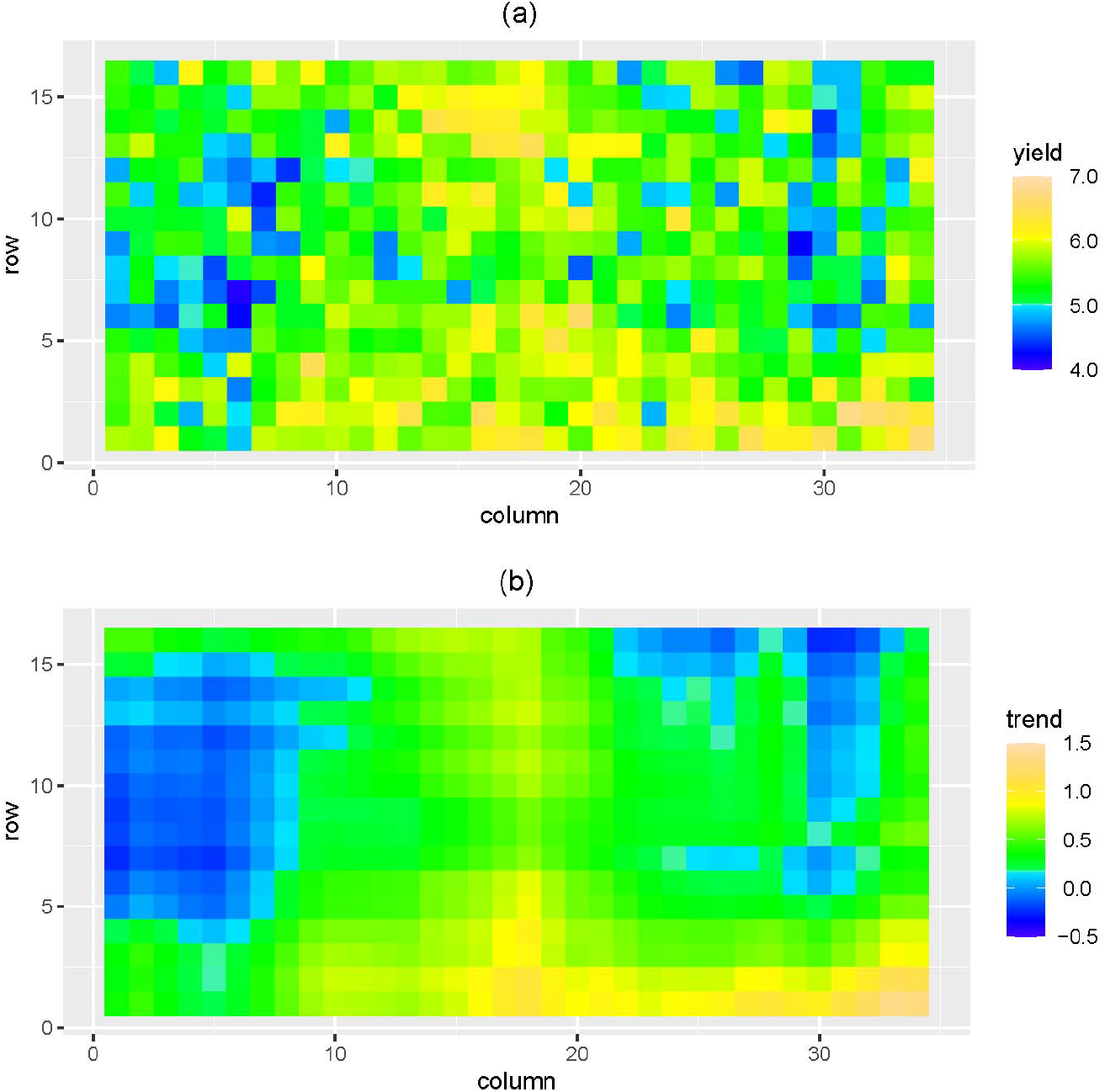
Heatmap of barley data of Durbán et al. (2003). (a) Raw data; (b) Smooth trend fitted by model M26 (Table 2).

Table 2 shows several models fits for *p* = 2, *i_r_* = *k* and *i_c_* = *s*, i.e., interior knots are placed at the plots. Model M12 with *q* = 3 and diagonal structures (Ω) for both marginal smooths was used to perform a grid search determining rotation angles maximizing the deviance approximately (Figure 2). These values were then used to fit the marginal smooths with unstructured covariance (∑). We considered *q* = 3 (cubic splines) and *q* = 1 (ordinary second differences; Green et al., 1985). A few observations are as follows:

i. The unstructured model is confirmed to ensure translation invariance, as opposed to the models with diagonal or constant variance. This can be seen by comparing models M3-M6 to M6-M8, where only *u_r_* is fitted. The diagonal models M6 and M8 fitted quite well compared to the invariant unstructured models (M3-M5). When also fitting *u_c_*, we were not able to fit models M16-M17 with unstructured ∑_*r*_ and ∑_*c*_ because positive definiteness of these matrices could not be enforced during iterations. Fixing the covariance in ∑*_c_* at zero, convergence was achieved. Note, however, that M16-M17 are equivalent to M15, which we were able to fit and which fitted best among the models in Table 2.
ii. M20 could not be fitted with *q* = 3 (again fixing the covariance in ∑_*c*_ at zero allowed convergence), but fitted second best with *q* = 1, giving a fit that was almost identical to M15.
iii. Using the 1*_k_* ⋮ *h_k_* as the representation of *X*_*r*_ in the marginal smooth gave poorer fits and also was more difficult to get to converge.
iv. There is only a very small difference between fits for *q* = 1 and *q* = 3, suggesting that the B-splines with *q* > 1 are not strictly needed and we can revert to simple first or second differences (Eilers, 2003) as in classical NN (Besag and Kempton, 1986; Williams, 1986; Wilkinson et al., 1983).

**Table 2:**
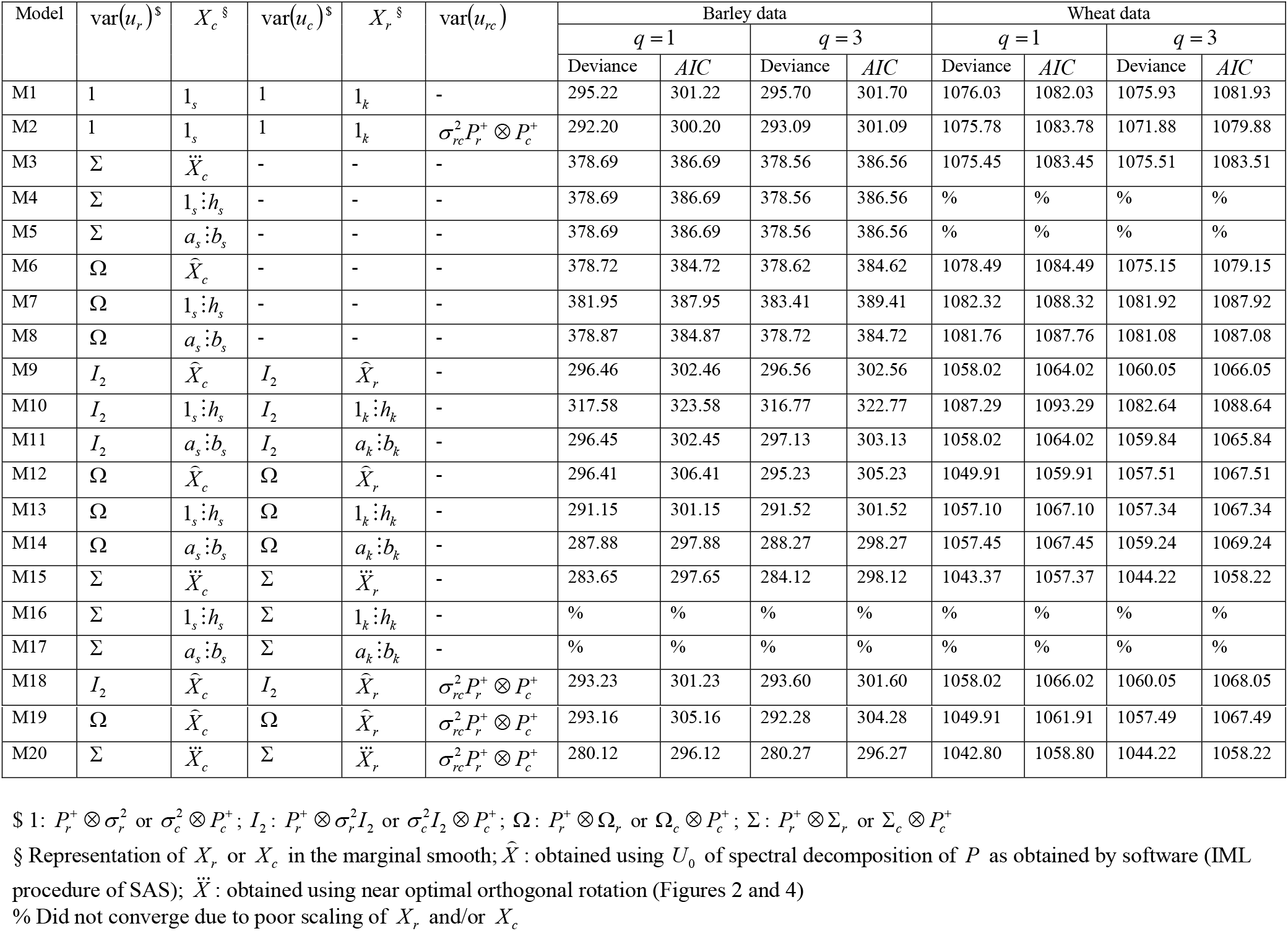
Analysis of barley data of Durbán et al. (2003) and wheat data of Stroup et al. (1994) using P-spline approach with *p* = 2 (second differences), *i_r_* = *k*, *i_c_* = *s*, and the penalty in (3) for *B_rc_*. All models have fixed effects for replicates, genotypes, row numbers *h_r_*, column numbers *h_c_* and their product *h_r_h_c_*.

**Figure 2:**
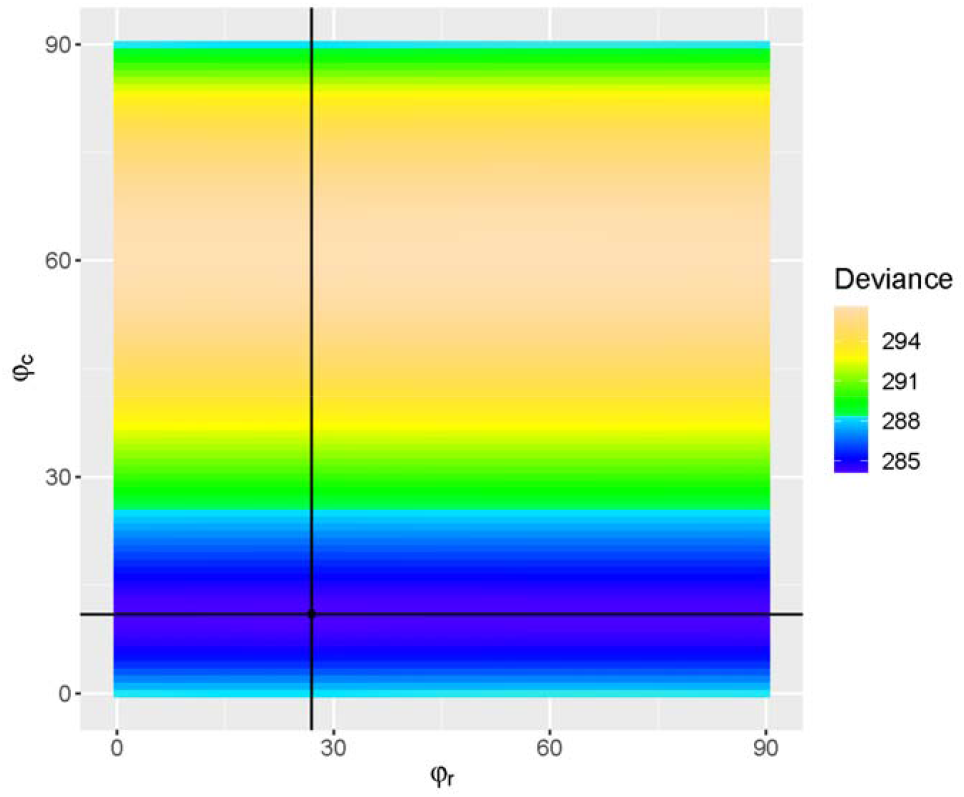
Deviance profiled for *φ_r_* and *φ_c_* for barley data using model M12 with *q* = 3 and diagonal structures (Ω) for both marginal smooths (Table 2). Minimum at *φ_r_* = 27° and *φ_c_* = 11°.

Table 3 shows results for first differences *p* = 1. These are quite competitive with second differences (*p* = 2). Reducing the number of knots had hardly any effect. The best fit is obtained for *q* = 1 and knots placed at the plots (M26) [see Figure 1(b) for the smoothed trend]. Table 4 shows the fits obtained with the IAR model with penalty (7) (Section 4.2) for a step size of 0.1 for *α* in the grid search. Again, first differences seemed to do better than second differences. Model M41 with *q* = 1 fitted best among all models considered for this dataset. For *p* = 2, we also fitted smooth marginal terms involving the full *X*_*r*_ and *X_c_*, though this would not be necessary to represent the null space of the interactions smooth. We faced the same problems when trying to fit the marginal penalty for columns with unstructured ∑*_c_*. When fixing the covariance to zero, convergence was easy to achieve. With the diagonal models for the marginal smooths, convergence was also unproblematic. For this model, we also report the fit for fixed *α* = 0.5 (M45 and M46), corresponding to the penalty in (9), for the case *q* = 3. This model is the one used in Rodríguez-Álvarez et al. (2018) and implemented in the SpATS R package, available from CRAN (https://cran.r-project.org/package=SpATS). The model is doing quite well in terms of AIC, but several others reported in Tables 3 and 4 fare slightly better. The best model was often obtained for *α* = 1.0, in which case the null space is not fully represented by the model; these models would need to be extended to fully cover the null space, which we have not pursued here. We also fitted the LV⊗LV model (Piepho and Williams, 2010), which is linear in the parameters, and the AR1⊗AR1 model, which is nonlinear (Gilmour et al., 1997) (Table 5). Both models also have good fits, with an edge in favour of the latter when a nugget was added. That model (M56) had a very similar fit to the best P-spline model in Table 3 (M26) but was inferior to the best model in Table 4 (M41).

**Table 3:** Analysis of barley data of Durbán et al. (2003) and wheat data of Stroup et al. (1994) using P-spline approach with *p* = 1 (first differences) and the penalty in (3) for *B_rc_*. All models have fixed effects for replicates, genotypes, row numbers *h_r_*, column numbers *h_c_* and their product *h_r_h_c_*. The marginal smooths use 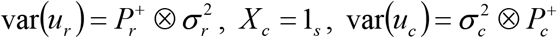 and *X_r_* = 1*_k_* for all models.

| Model | $q$ | $i_r$ | $i_c$ | $\text{var}(u_{rc})$ | Barley data | | Wheat data | |
| --- | --- | --- | --- | --- | --- | --- | --- | --- |
|  |  |  |  |  | Deviance | AIC | Deviance | AIC |
| M21 | 3 | $k$ | $s$ | - | 293.37 | 299.37 | 1072.47 | 1078.47 |
| M22 | 3 | $k$ | $s$ | $\sigma_{rc}^2 P_r^+ \otimes P_c^+$ | 279.28 | 287.28 | 1046.40 | 1054.40 |
| M23 | 2 | $k$ | $s$ | - | 293.56 | 299.56 | 1071.91 | 1077.91 |
| M24 | 2 | $k$ | $s$ | $\sigma_{rc}^2 P_r^+ \otimes P_c^+$ | 279.18 | 287.18 | 1046.06 | 1054.06 |
| M25 | 1 | $k$ | $s$ | - | 295.78 | 301.78 | 1075.14 | 1081.14 |
| M26 | 1 | $k$ | $s$ | $\sigma_{rc}^2 P_r^+ \otimes P_c^+$ | 278.45 | 286.45 | 1047.12 | 1055.12 |
| M27 | 3 | 10 | 20 | - | 292.92 | 298.92 | 1073.38 | 1079.38 |
| M28 | 3 | 10 | 20 | $\sigma_{rc}^2 P_r^+ \otimes P_c^+$ | 281.18 | 289.18 | 1056.41 | 1064.41 |
| M29 | 2 | 10 | 20 | - | 291.61 | 297.61 | 1073.27 | 1079.27 |
| M30 | 2 | 10 | 20 | $\sigma_{rc}^2 P_r^+ \otimes P_c^+$ | 279.74 | 287.74 | 1057.09 | 1065.09 |
| M31 | 1 | 10 | 20 | - | 296.75 | 302.75 | 1075.43 | 1081.43 |
| M32 | 1 | 10 | 20 | $\sigma_{rc}^2 P_r^+ \otimes P_c^+$ | 281.31 | 289.31 | 1052.41 | 1060.41 |

**Table 4:** Analysis of barley data of Durbán et al. (2003) and wheat data of Stroup et al. (1994) using the IAR model with the penalty in (7) for *u_rc_* with *i_r_* = *k* and *i_c_* = *s* with and without marginal smooths added. All models have fixed effects for replicates, genotypes, row numbers *h*_*r*_, column numbers *h_c_* and their product *h_r_h_c_*. Fits obtained by a grid search over *α* = 0,1(0.1).

| Model | $q$ | $p$ | Marginal smooth<sup>§</sup> | Barley data | | | Wheat data | | |
| --- | --- | --- | --- | --- | --- | --- | --- | --- | --- |
| | | | | $\alpha$ | Deviance | AIC | $\alpha$ | Deviance | AIC |
| M33 | 3 | 1 | - | 0.5 | 284.53 | 290.53 | 0.9 | 1052.33 | 1058.33 |
| M34 | 2 | 1 | - | 0.5 | 283.94 | 289.94 | 0.9 | 1053.18 | 1059.18 |
| M35 | 1 | 1 | - | 0.6 | 281.52 | 287.52 | 0.8 | 1055.06 | 1061.06 |
| M36 | 3 | 2 | - | 0.6 | 295.90 | 301.90 | 0.9 | 1063.35 | 1069.35 |
| M37 | 2 | 2 | - | 0.6 | 296.58 | 302.58 | 0.9 | 1064.66 | 1070.66 |
| M38 | 1 | 2 | - | 0.6 | 297.00 | 303.00 | 0.9 | 1064.94 | 1070.94 |
| M39 | 3 | 1 | 1 | 0.9 | 274.84 | 284.84 | 0.9 | 1050.84 | 1058.84 |
| M40 | 2 | 1 | 1 | 0.9 | 274.85 | 284.85 | 1.0 | 1050.61 | 1060.61 |
| M41 | 1 | 1 | 1 | 0.9 | 273.81 | 283.81 | 0.9 | 1053.08 | 1063.08 |
| M42 | 3 | 2 | 1 | 1.0 | 284.63 | 294.63 | 0.9 | 1059.55 | 1069.55 |
| M43 | 2 | 2 | 1 | 1.0 | 284.79 | 294.79 | 0.9 | 1060.54 | 1070.54 |
| M44 | 1 | 2 | 1 | 1.0 | 284.94 | 294.94 | 0.9 | 1061.39 | 1071.39 |
| M45<sup>%</sup> | 3 | 2 | $\Omega$ | 0.5 | 283.58 | 295.58 | 0.5 | 1063.57 | 1077.57 |
| M46<sup>§</sup> | 3 | 2 | $\Omega$ | 0.5 | 290.92 | 302.92 | 0.5 | 1052.70 | 1066.70 |
| M47 | 3 | 2 | $\Omega$ | 1.0 | 285.24 | 297.24 | 1.0 | 1049.40 | 1063.40 |
| M48 | 2 | 2 | $\Omega$ | 1.0 | 287.06 | 299.06 | 1.0 | 1044.35 | 1058.35 |
| M49 | 1 | 2 | $\Omega$ | 1.0 | 287.20 | 299.20 | 1.0 | 1046.99 | 1060.99 |
| M50 | 3 | 2 | $\Sigma$ | 1.0 | 273.67 | 289.67 | 1.0 | 1044.28 | 1062.28 |
| M51 | 2 | 2 | $\Sigma$ | 1.0 | 274.77 | 290.77 | 1.0 | 1041.51 | 1059.51 |
| M52 | 1 | 2 | $\Sigma$ | 1.0 | 273.51 | 289.51 | 1.0 | 1038.81 | 1056.81 |

§ 1: for $p = 1$ just the intercepts for rows and columns. $\Omega$ : for $p = 2$ using $P_r^+ \otimes \Omega_r$ and $\Omega_c \otimes P_c^+$ with $\hat{X}_c$ and $\hat{X}_r$ , respectively. $\Sigma$ : using $P_r^+ \otimes \Sigma_r$ and $\Sigma_c \otimes P_c^+$ with $\ddot{X}_c$ and $\ddot{X}_r$ , respectively; $\hat{X}$ : obtained using $U_0$ of spectral decomposition of $P$ as obtained by software (IML procedure of SAS); $\ddot{X}$ : obtained using near optimal orthogonal rotation (Figures 2 and 4)
% Fit for fixed $\alpha = 0.5$ ; using $\hat{X}_c$ and $\hat{X}_r$ for $X_c$ and $X_r$ , respectively.
§ Fit for fixed $\alpha = 0.5$ ; using $a_s:b_s$ and $a_k:b_k$ for $X_c$ and $X_r$ , respectively (Rodríguez-Álvarez et al. 2018).

**Table 5:**
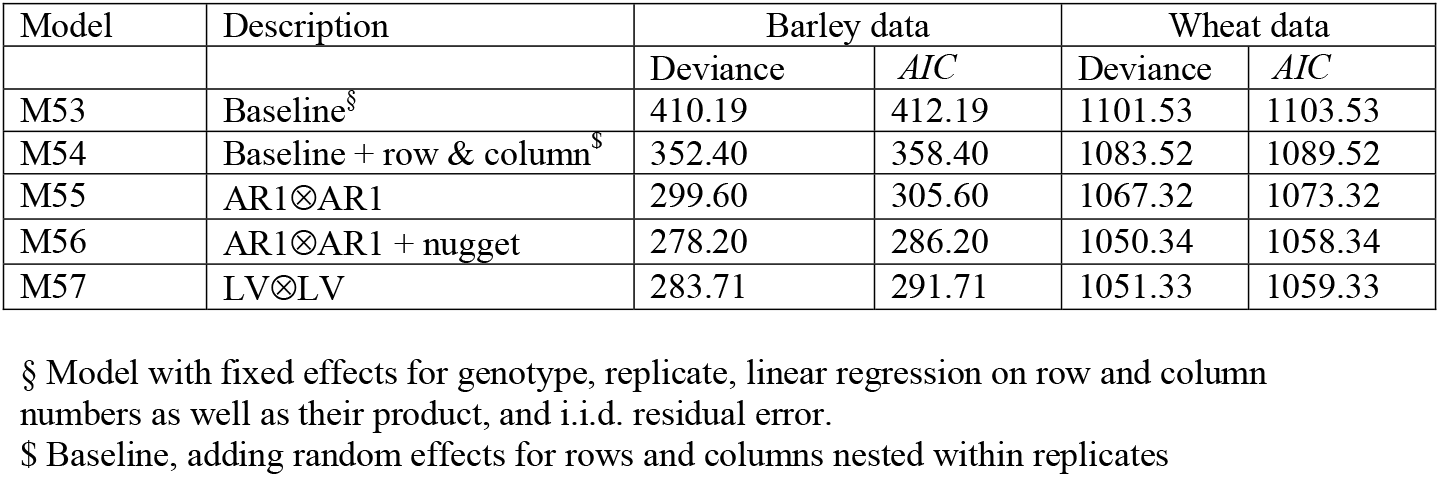
Analysis of barley data of Durbán et al. (2003) and wheat data of Stroup et al. (1994) using other common models. All models have fixed effects for replicates, genotypes, row numbers *h_r_*, column numbers *h_c_* and their product *h_r_h_c_*.

### 5.2. A wheat trial

Stroup et al. (1994) report a trial for 56 wheat varieties laid out in *k* = 11 rows and *s* = 22 (Figure 3) (this is the ‘Alliance’ trial in their paper). The dataset displays a strong spatial trend, which is described by Stroup (2002) in these terms: ‘The pattern is typical of spatial variability. In this case, relatively low yields tend to be clustered in the northwest corner. This pattern is explained by a low rise in this part of the field causing increased exposure to winter kill from wind damage, and hence depressed yield.’ We here use the version of the dataset that is available in the ‘agridat’ package of R (Wright, 2021; https://CRAN.R-project.org/package=agridat), which has the yield expressed in bushels per acre. The four replicates are not arranged as a rectangular array, with some rows divided up between two replicates. We fitted the exact same models as for the barley data (Tables 2 to 5). The general picture emerging from this second example is rather similar to that from the first, so details will not be dwelt on here. It is worth mentioning that M46, which corresponds to SpATS (Rodríguez-Álvarez et al,. 2018), gives a good fit in terms of AIC, landing in the mid-range of the other models reported in Tables 3 and 4. The unstructured models for the marginal smooths could only be fitted with the rotation determined by a two-dimensional grid search for the diagonal model (Figure 4). Even with this rotation, it was not easy to achieve convergence, mainly due to a variance component approaching zero. All other models were easy to fit. The results for *p* = 2 do confirm that the diagonal model is not invariant to rotations but the unstructured model is. One main conclusion from this example is that first differences work quite well.

**Figure 3:**
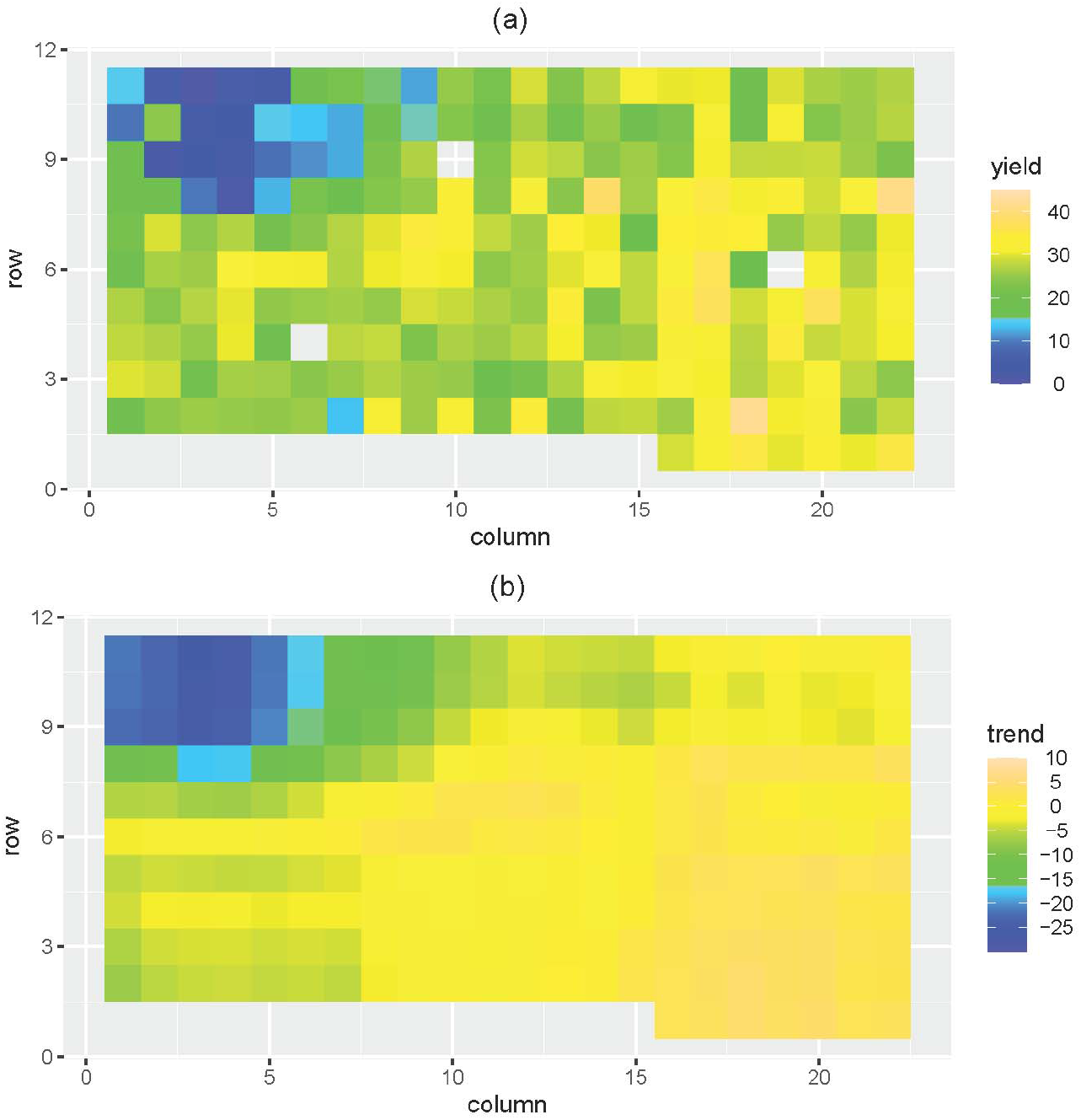
Heatmap of wheat data of Stroup et al. (1994). (a) Raw data; (b) Smooth trend fitted by model M26 (Table 2).

**Figure 4:**
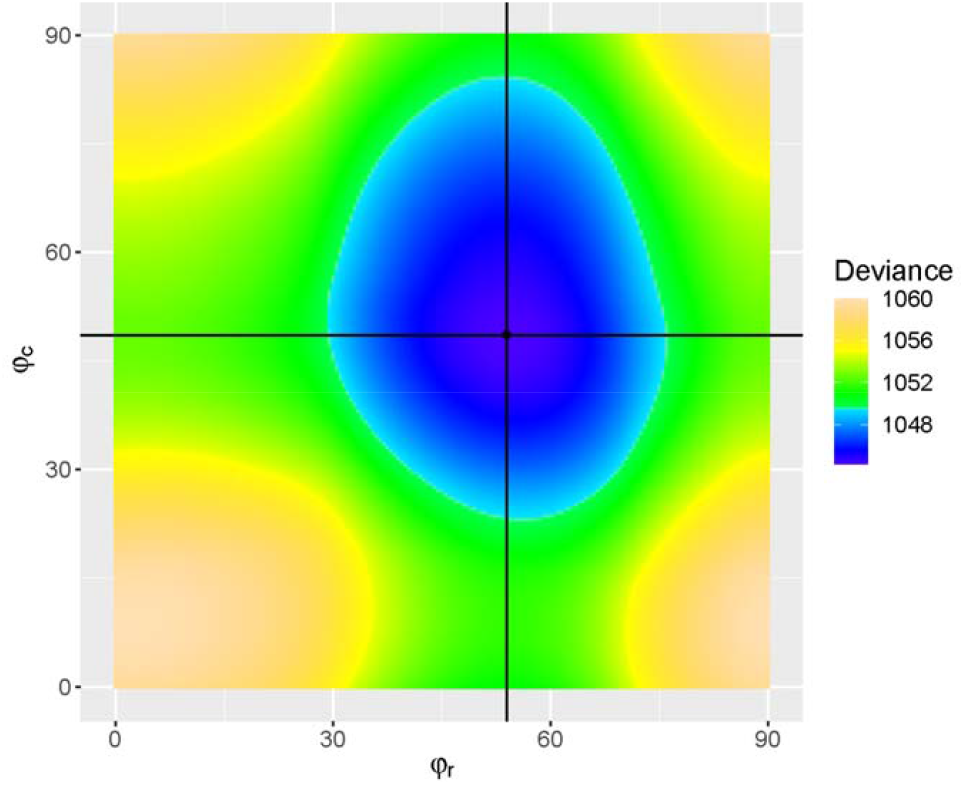
Deviance profiled for *φ_r_* and *φ_c_* for wheat data using model M12 (Table 2). Minimum at *φ_r_* = 52° and *φ_c_* = 48°.

## 6. Discussion

We considered a framework that allows the use of P-splines to model spatial trends in field trials (Rodríguez-Álvarez et al., 2018). The methodology requires making several choices, i.e., the order of differencing (*p*), the degree of the B-spline basis (*q*), and the number of knots (*i_r_*, *i_c_*), which in field trials are always placed on a regular grid. As the two examples illustrate, there is indeed a very large number of options, and this means that model choice could easily become an overwhelming exercise. It would be advantageous for routine use if a limited set of models could be identified that works well over a broad range of settings, not requiring an ‘intricate process of tailoring models to individual datasets’ which always entails ‘an element of subjectivity which will be difficult to eliminate’ (Rob Kempton, in discussion of Verbyla et al., 1999). A natural starting point for the placement of knots are the plot locations, and this is our general recommendation. Our results for the barley and wheat data suggest that there is little gain from trying other options. We also found little difference in goodness-of-fit for different choices of the degree of the B-spline basis, with the simplest choice of *q* = 1 (implying the simplification *B* = *I*) working quite well, corresponding to the classical NN modelling with first or second differences for *p* = 1 and *p* = 2, respectively. The examples further showed that first differences seemed to be sufficient for smoothing the spatial trend. Certainly, more empirical experience is needed to further provide guidance for routine use of the framework. We do conjecture based on our experience so far that a viable recommendation is to stick with the most parsimonious options, such as *p* = 1, *q* = 1, *i_r_* = *k*, and *i_c_* = *s*. These models are particularly easy to fit. This option is also very closely related to earlier approaches to NN analysis of field trials and so will be easy to communicate to researchers wanting to use P-splines. For practical implementation, one may either use the specialized SpATS R package, which implements a subset of the models considered in this paper, or a general linear mixed model package such as SAS, for which sample code is provided in the supplemental material.

Careful consideration needs to be given to the null space of the interaction smooth, especially for the models described in Section 4.1. As detailed there, this null space can have a rather larger number of dimensions for *p* = 2 than for *p* = 1 when the penalty in (3) is used. If one insists on representing this null space by fixed effects in order to make the fit invariant to the choice of g-inverse for the singular precision matrix *P_rc_*, efficiency of the analysis may be adversely affected, and it is therefore worthwhile to fit these effects as random. In particular, the P-spline framework allows smoothing these terms, combining them with the marginal smooth for rows and columns as prescribed by the ANOVA-type approach to spline smoothing (Wood et al., 2013; Wood, 2017). It must be acknowledged, however, that this sacrifices the invariance to the choice of g-inverse for *P_rc_*. We note in this context that the LV⊗LV model gives the same fit as the P-spline approach for *p* = 1, *q* = 1, *i_r_* = *k*, and *i_c_* = *s*, when the null space is modelled by fixed effects for rows and columns, but not when these effects are modelled as random (Boer et al., 2020), or are smoothed (Tables 3 and 5). An advantage of the IAR model with the penalty in (7) is the small size of the null space for the interaction smooth, even for *p* = 2, with a simple fixed-effects regression on row and column numbers to cover that null space. Thus, there is no confounding with the marginal smooths and so these can be considered in their own right, there being no strict need to fit them. Care is needed with the interaction smooth, however, when *α* converges to either 0 or 1 (this happened with both examples), because in that event the null space changes, increasing considerably, which may be seen as a disadvantage of the IAR model. One way out is to constrain the grid search so that *α* stays clear of the lower and upper bound, or use specialized procedures such as the SAP algorithm outside standard mixed model packages (Rodríguez-Álvarez, 2015), and implemented in the SpATS R-package.

We have included fixed-effects regression terms for row and column numbers and their interaction in all models, even those that do not involve P-splines, and also for P-splines with first-difference penalty, where these terms are not needed to represent the null space of the smooth. This was done so we could use the REML-based likelihood for comparing all models on the same basis. These terms only cost three degrees of freedom in the fixed part of the model, and it may therefore be worth including them in their own right in order to capture global trends, even if they are not needed to represent the null space of the smooth terms.

A general advantage of the P-spline framework is that most of the variance-covariance structures are linear in the parameters, which makes them easy to fit and minimizes convergence problems which may befit spatial variance-covariance models that are nonlinear in the parameters (Piepho et al., 2015; Velazco et al., 2017). This plays out most when the random terms are all independent, i.e. the smooth terms correspond to a variance-components model with no covariances among random effects. As we have shown, when *p* > 1, the marginal smooth associated with the penalty (3) for the interaction may require allowing a covariance among random intercept and slope terms, a key result that seems to have been overlooked by others, and this can be difficult to fit on account of the need to ensure positive definiteness of the variance-covariance structure. Such difficulties pose an obstacle for routine use, and they speak in favour of the simpler P-spline options based on first differences. In this regard, it is interesting to observe that the AR1⊗AR1 model with a nugget variance (M56), which fitted quite well, had relatively large correlations along rows and columns (0.8598 and 0.8952 for the barley data and 0.7488 and 0.9064 for the wheat data), in which case the covariance is well approximated by a linear variance model (Williams, 1986), corresponding to the simpler P-spline for *p* = 1, *q* = 1, *i_r_* = *k*, and *i_c_* = *s*.

On a final note we would like to stress the importance of design, which often does not receive the attention it deserves. Sometimes, the large number of modelling options for spatial analysis may raise the false impression that design does not matter, and that a sophisticated analysis takes care of everything. Nothing could be further from the truth. Certainly, given the large number of options for analysis, the design issue becomes more challenging and is far from resolved. Rather than focusing the design on a specific route of analysis, one may consider searching for a good design without a specific analysis in mind (Piepho et al., 2021), striving for robustness to model choice.

## CRediT authorship contribution statement

**Hans-Peter Piepho:** Conceptualization, Methodology, Formal analysis, Writing - original draft. **Martin P. Boer:** Conceptualization, Methodology, Formal analysis, Writing - review and editing. **Emlyn R. Williams:** Conceptualization, Methodology, Formal analysis, Writing - review and editing.

## Appendices

### A. Row-wise Kronecker products

Eilers and Marx (2003) proposed the use of row-wise Kronecker products, also known as box products or tensor products of B-spline bases for non-additive smoothing. A salient feature of field trials, however, is that plots are usually placed on a regular grid, for which the tensor product coincides with the usual Kronecker product (Lee and Durbán, 2011). Thus, if ⊗*_r_* denotes the row-wise Kronecker product and ⊗ denotes the ordinary Kronecker product, then for a B-spline basis matrix *B_r_* with *k* rows and arbitrary number of columns corresponding to regularly spaced knots and a B-spline basis matrix *B_c_* with *s* rows and an arbitrary number of columns (knots), we have

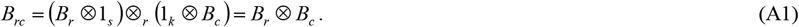

This important fact can hardly be over-emphasized for those wanting to use P-splines for analysis of spatial data on a regular grid in general and on a regular grid of field trial plots in particular, but drawing on a literature that seeks to embrace irregularly gridded data as the dominant application case, e.g. in longitudinal studies. In the case of field trials this means that a row-wise Kronecker product can be written as an ordinary Kronecker product of matrices associated with rows and columns (Verbyla et al., 2018), which greatly simplifies the notation and also clarifies the relationship with other spatial modelling approaches for field trials, such as the LV⊗LV model (Williams and Piepho, 2010) and the AR1⊗AR1 model (Gilmour et al., 1997). We make use of the important equation (A1) as a design matrix throughout this paper to model spatial interactions between rows and columns. Readers less familiar with P-splines should be aware that the simplified notation arising from (A1) may look somewhat unfamiliar compared to results for P-splines applicable to irregular spatial grids (Lee and Durbán, 2011; Wood, 2017, p. 232), even though the regular case is just a special version of the irregular one.

### B. Relations with state-space models

The use of differences among neighbouring plots (Besag and Kempton, 1986) has close ties with state-space models (Harvey, 1989; Durbin and Koopman, 2001), which in turn can also be represented as mixed models (Tsimikas and Ledolter, 1997; Piepho and Ogutu, 2007). Specifically, we may assume that the response *y_i_* on the *i*-th plot is related to a latent level *μ_i_* according to *y_i_* = *μ_i_* + *e_i_*, where 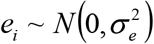. If we further assume that the state on the neighbouring (*i*+1)-th plot is related to that on the *i*-th plot by *μ*_*i*+1_ = *μ_i_*+ *u_i_*, where 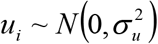, then *μ_i_* follows a *first-order random walk* model, which is a simple form of state-space model. For first differences we then have 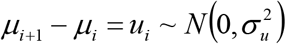, demonstrating the connection between the use of first differences and state-space models as used mostly for time series. By comparison, the AR1 model is known as damped model in state-space modelling and written as *μ*_*i*+1_ = *ρμ_i_* + *u_i_* with 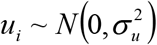, where 0 < *ρ* < 1 is the autocorrelation (Harvey, 1989, p. 46).

The first-order random walk model can be extended to what is known in state-space modelling as *local linear trend* model. The starting point is a simple linear regression of the form *μ_i_* = *β*_0_+ *β*_1_*i*. One idea to make this deterministic trend stochastic is to let both *β*_0_ and *β*_1_ vary randomly, but this would lead to discontinuities at the plots (Harvey, 1989, p. 37). A better option is to work directly with the current level, *μ_i_* in place of the intercept *β*_0_. Noting that the deterministic trend can be rewritten as *μ*_*i*+1_ = *μ_i_*+ *β*_1_ with *μ*_0_ = *β*_0_, we may consider updating both the current level and the slope term by a random walk as follows:

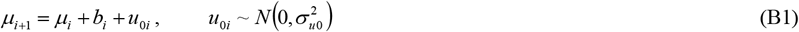

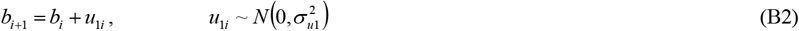

where *b_i_* is the random slope at the *i*-th plot with *b*_0_ = *β*_1_. Thus, on the *i*-th plot we have linear trend *b_i_*, and this is updated by a random increment *u*_1*i*_, i.e. by a random walk, as we move to the (*i*+1)-th plot. The intercept at the *i*-th plot is *a_i_* = *μ_i_* – *b_i_i*, and hence *a*_*i*+1_ = *μ_i_* – (1 – *i*)*b_i_* + *u*_0*i*_. Essentially, we have randomly varying regression lines, pieced together at the plots. Due to the update *u*_0*i*_, however, there are discontinuities at the plots (Figure 1). In the limiting case when 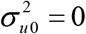, the trend is a polygon with breaks at the plots but no discontinuities. Interestingly, the model then is equivalent to *μ*_*i*+1_ – 2*μ*_*i*_ + *μ*_*i*–1_ = *u*_1*i*_, which is a *second-order random walk* and involves second differences (Durbin and Koopman, 2001, p. 39). To see this, we may subtract the state equation 0 = – *μ_i_* + *μ*_*i*–1_ + *b*_*i*–1_ from *μ*_*i*+1_ = *μ_i_* + *b_i_*, yielding 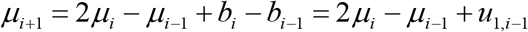. By comparison, the first-order random walk model for *μ_i_* can be depicted as horizontal regression lines with jumps at the plots (Figure B1). It is worth observing that both the local linear trend and its limiting case with 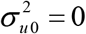, which is equivalent to use of second differences, have the fixed trend component *β*_0_ + *β*_1_*i*, corresponding to the null space of P-splines with second differences penalty. Thus, the state-space view provides a very natural explanation why a fixed intercept as well as a fixed slope are needed with second differences.

**Figure B1:**
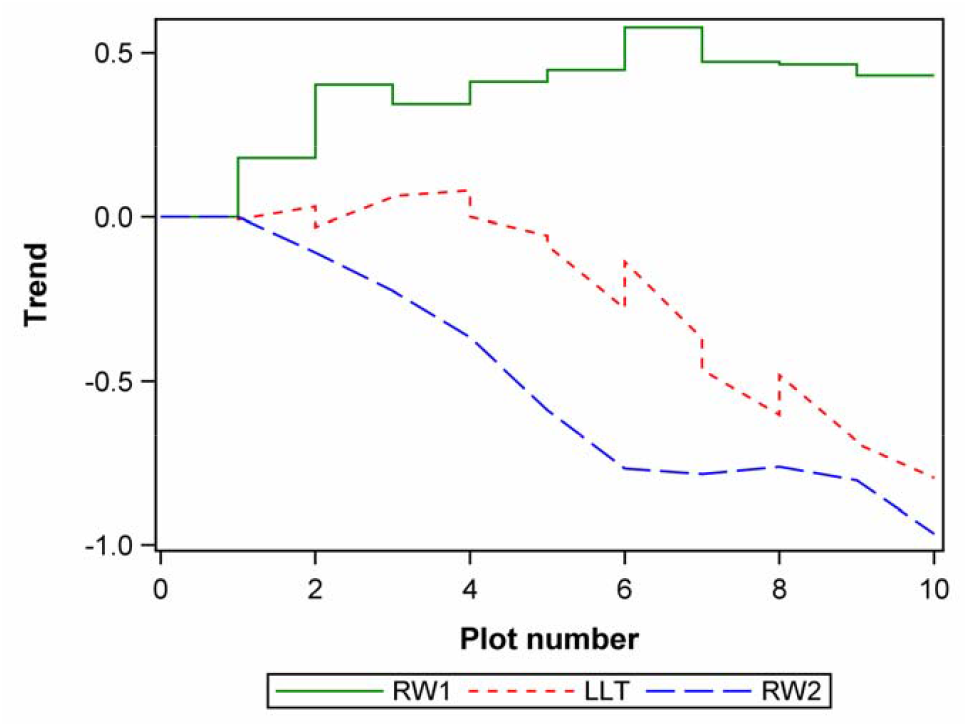
Simulated trend for three state-space models. RW1 = first-order random walk; RW2 = second-order random walk; LLT = local linear trend; *μ*_0_ = *b*_0_ = 0; 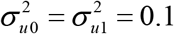.

### C. Two-dimensional first differences

Kempton et al. (1994) formulate a correlation structure that is separable between rows and columns. Their assumption is that first differences along rows and along variances have variances *ϕ_r_* = *ψ_r_σ*^2^ and *ϕ_c_* = *ψ_c_σ*^2^ respectively, and that plot differences calculated across both rows and column, i.e., *v_i,j_* – *v*_*i*+1,*j*_ – *v*_*j,i*+1,_ + *v*_*i*+1,*j*+1_, where *v_i,j_* – *v*_*i*+1,*j*_ – *v*_*j,i*+1,_ + *v*_*i*+1,*j*+1_ is the plot value in the *i*-th row and *j-th* column, have variance *ϕ_rc_* = (2 +*ψ_r_*)(2 + *ψ_c_*)*σ*^2^. In what follows, we will relate this model to the LV⊗LV model (Piepho and Williams, 2010) and P-splines with *p* = 1, *q* = 1, *i_r_* = *k*, and *i_c_* = *s*. We may collect plot values for spatial trend into a random vector *v_rc_*. Following Kempton et al. (1994), the first differences across rows can then be assumed to have variance

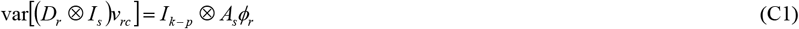

for some *s* × *s* correlation matrix *A_s_*. For first differences across columns, we may postulate accordingly that

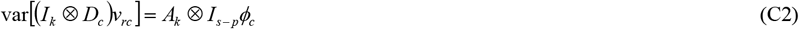

for some *k* × *k* correlation matrix *A_k_*, and for differences across both rows and columns

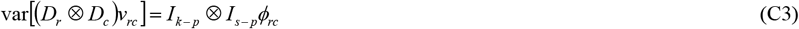

In line with our P-spline approach, the variance-covariance matrix for *v_rc_* may be assumed to take the form

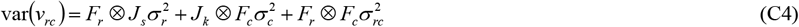

for suitable variance-covariance matrices *F_r_* and *F_c_*. This can also be expressed in terms an ANOVA-type decomposition

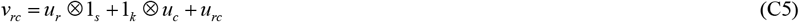

with 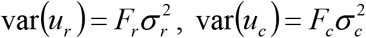 and 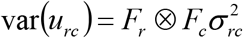. We now need to find *F_r_* and *F_c_* such that 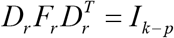 and 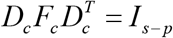. For *F_r_*, e.g., we can meet these assumptions using 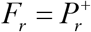 or 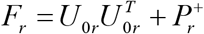, corresponding to our P-spline approach, but this does not yield the desired form of *A_s_*. One (but not the only) choice that will yield the desired form of *A_s_* is 2*F_r_* = *M_k_* = *J_k_*(*k* –1)–*L_k_*, where *L_k_* is an *k* × *k* matrix with (*i*_1_, *i*_2_)-th element equal to |*i*_1_ – *i*_2_|, corresponding to the LV⊗LV model (Piepho and Williams, 2010). To see this, we first use (C4) to find

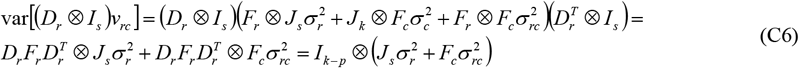

Comparing coefficients with (C1), we find 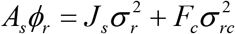. Then using 2*F_r_* = *M_k_*, we find that the diagonal elements of *A_s_ϕ_r_* equal 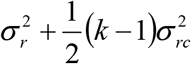, and hence 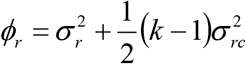. Similarly, 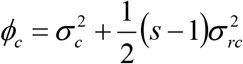. Moreover, 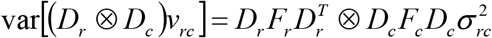, from which 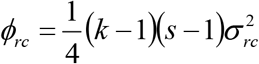.

### D. Orthogonal rotations of the null-space eigenvectors for p = 2

It is interesting to observe the connection of a diagonal variance-covariance model with rotation parameter with an unstructured model. Let Ω be a diagonal matrix

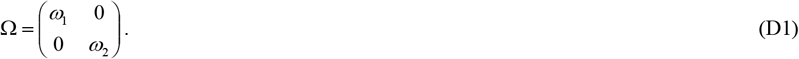

Then using *U*_0_ = (*a_m_*⋮*b_m_*)*R*(*φ*) [see eqs. (5) and (6)] we have

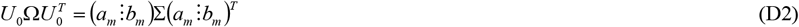

where

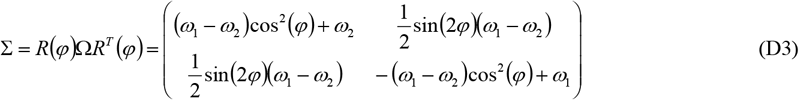

If *ω*_1_ = *ω*_2_ = *ω*, we have Σ = Ω = *ωI*_2_, showing that if we chose the identity matrix, the eigenvectors can be chosen arbitrarily, i.e. the model does not depend on the rotation parameter *φ*.

## Notes

### Competing Interest Statement

The authors have declared no competing interest.

